# P2Y2 purinergic receptor and DNA sensor cGAS dictate ionizing radiation-mediated proinflammatory macrophage activation

**DOI:** 10.1101/2024.12.19.629560

**Authors:** Ali Mostefa-Kara, Aurélia Deva Nathan, Audrey Paoletti, Anais Chapel, Emie Gutierrez-Mateyron, Thanh Trang Cao-Pham, Yara Bou Ghosn, Awatef Allouch, Jean-Luc Perfettini

## Abstract

The reprogramming of tumor-associated macrophages (TAMs) by radiotherapy is associated with cancer patient’s response and sensitization to immune checkpoint blockade, but the molecular mechanisms involved remain largely unknown. Here, we show that following ionizing radiation (IR), macrophages accumulate single and double strand-DNA breaks and fragmented mitochondria in their cytosol, and stabilize the DNA sensor cyclic GMP-AMP synthase (cGAS). We demonstrate that mitochondrial fragmentation is induced by the activation of the dynamin-related protein 1 (DRP1), and controls the stabilization of cGAS and the proinflammatory activation of irradiated macrophages. Furthermore, pharmacological and genetic inhibitions of cGAS impair the proinflammatory activation of irradiated macrophages, thus revealing that cGAS is a central effector of IR-mediated proinflammatory macrophage activation. Interestingly, we also report that the purinergic receptor P2Y2 acts as an endogenous repressor of the proinflammatory macrophage activation and demonstrate that P2Y2 inactivation enhances the capacity of irradiated macrophages to undergo a proinflammatory activation. Our results thus define a new signaling pathway elicited in macrophages by IR directing mitochondrial dynamics, cytosolic DNA recognition by cGAS and proinflammatory phenotype, which is enhanced following P2Y2 inactivation.

## Introduction

Every year, approximately thirty to fifty percent of cancer patients are treated worldwide with radiotherapy alone or in combination with other cancer treatments such chemotherapies and surgery [1]. Despite the central place of radiotherapy in the therapeutic armamentarium available to clinicians for cancer treatment, the cellular and molecular mechanisms responsible for the efficacy of radiotherapy are partially understood. In response to IR, cancer cells can be directly eliminated through the induction of distinct cell-autonomous death (CAD) or non-cell-autonomous death (NCAD) modalities [2] (including apoptosis [3], autophagic cell death [4], necroptosis [5], methuosis [6], cellular cannibalism [7], ferroptosis [8] and mitotic catastrophe [9]) or senescence [10]. These cytotoxic processes can be associated with the release of immunogenic damage-associated molecular patterns (DAMPs) (such as adenosine triphosphate (ATP) [11] and high mobility group box 1 (HMGB1) [12]) and the induction of both intracellular and extracellular immune-stimulatory signals (such as cytosolic single- or double-strand DNA breaks [13] and type I Interferon β (IFN-β) release [14]) that support the development of an effective antitumor immune response. IR can also modulate functions of non-cancerous cells of the tumor microenvironment (TME) (including endothelial cells, cancer-associated fibroblast and immune cells), reshape the architecture of TME and trigger associated immune-modulatory effects [15, 16].

TAMs are major cellular components of TME that sustain in the vast majority of cancers, tumor immune evasion through anti-inflammatory activation [17] and tumor phagocytosis defectiveness [18], thus promoting cancer progression and resistance to cancer treatments [19]. The abrogation of macrophage immunosuppressive activities with therapeutic approaches such as monoclonal antibody against CD47 [20], PI3Kγ inhibitor [21] or functionalized nanoparticles [22] reveals that the proinflammatory reprogramming of TAMs directs tumor response and better clinical outcomes. Radiotherapy was initially revealed for its ability to reshape TME through vasculature normalization and indirect proinflammatory reprogramming of TAMs [23, 24]. We demonstrated that IR also directly acts on TAMs and induces their functional reprogramming from anti-inflammatory to proinflammatory phenotype [25]. This cell-autonomous process implies the activation of the NADPH oxidase 2 (NOX2), the ataxia telangiectasia mutated (ATM) kinase and the interferon related factor 5 (IRF5). The inhibition of this proinflammatory signaling is associated with the poor tumor response to preoperative adjuvant radiotherapy in locally advanced rectal cancer [25], thus supporting the idea that therapeutic proinflammatory reprogramming of TAMs should help to improve radiotherapy efficacy. Here, we further explored molecular and cellular mechanisms involved in the regulation of IR-mediated proinflammatory macrophage activation.

## Results

### Cytoplasmic DNA breaks and cGAS accumulate in irradiated macrophages during proinflammatory activation

Considering the capacity of IR to induce the release of genomic or mitochondrial DNA into the cytosol of irradiated cells [26, 27] and as a consequence, to regulate central macrophage functions such as antigen presentation and costimulatory molecule expression [28] and cytokine secretion [25], the accumulation of single-stranded (ss) and double-stranded (ds) DNA breaks in the cytosol of macrophages during IR-mediated proinflammatory activation was analyzed. Phorbol myristate acetate (PMA)-differentiated human THP1 macrophages were irradiated with a single dose of 2 Gy and analyzed using fluorescence microscopy. After 30 minutes, cytosolic puncta containing dsDNA breaks accumulated in approximately 32% of 2 Gy-irradiated PMA-differentiated human THP1 macrophages (Fig. 1A, B), as compared to control, non-irradiated PMA-differentiated human THP1 macrophages. Interestingly, ssDNA-containing puncta were also detected in the cytosol of 46% of 2 Gy-irradiated, PMA-differentiated human THP1 macrophages (Fig. 1C, D), underlining the fast accumulation of both ssDNA and dsDNA fragments in the cytosol of irradiated macrophages. To determine whether the accumulation of cytosolic DNA fragments could mirror the increased DNA damage detected in the nucleus of irradiated macrophages that we previously reported [25], the phosphorylation of the histone variant H2AX on serine 139, also known as γ- H2AX, was studied by fluorescence microscopy 1 hour after the irradiation of PMA-differentiated human THP1 macrophages with a single dose of 2 Gy. Approximately 50% of 2 Gy-irradiated PMA-differentiated human THP1 macrophages exhibited nuclear γ-H2AX^+^ foci (Supplementary Fig. S1A, B), as compared to control, non-irradiated PMA-differentiated human THP1 macrophages, thus revealing that the cytosolic accumulation of DNA fragments is associated with the induction of nuclear DNA damage in irradiated macrophages. The expression level of cGAS, which detects cytosolic DNA [29] and plays a central role in various biological processes elicited by IR, such as the secretion of type I interferon [14, 30], the induction of senescence [31] and the activation of antitumor immunity [13], was also determined using western blots. The increased expression levels of cGAS in murine RAW264.7 macrophages (Fig. 1E) or PMA-differentiated human THP1 macrophages (Fig. 1F), 3 hours and 6 hours after 2Gy irradiation were detected respectively, as compared to control, non-irradiated cells. Ninety-six hours after a single dose of 2 Gy, the increased expression levels of the inducible nitric oxide synthase (iNOS) and the interferon regulatory factor 5 (IRF5), which are two proinflammatory markers of macrophage activation [32, 33], were detected in PMA-differentiated human THP1 macrophages. As previously published [25], we confirmed using fluorescence microscopy that 24 hours after a single-dose of 2 Gy, the frequency of irradiated PMA-differentiated human THP1 macrophages expressing iNOS is significantly increased (Fig. 1G, H), as compared to control PMA-differentiated human THP1 macrophages. Accordingly, the increased expression level of IRF5 is also detected using western blots at 96 hours in 2 Gy-irradiated PMA-differentiated human THP1 macrophages (Fig. 1I), as compared to control PMA-differentiated THP1 macrophages. Of note, we detected the expression level of IRF5 after 6 hours in 2 Gy-irradiated murine RAW264.7 macrophages by western blots (Fig. 1J), thus showing that the increased expression level of cGAS occurs before the proinflammatory activation of irradiated macrophages. Altogether, these results reveal that cytosolic DNA breaks and cGAS accumulate in macrophages during IR-mediated proinflammatory activation.

**Fig 1:**
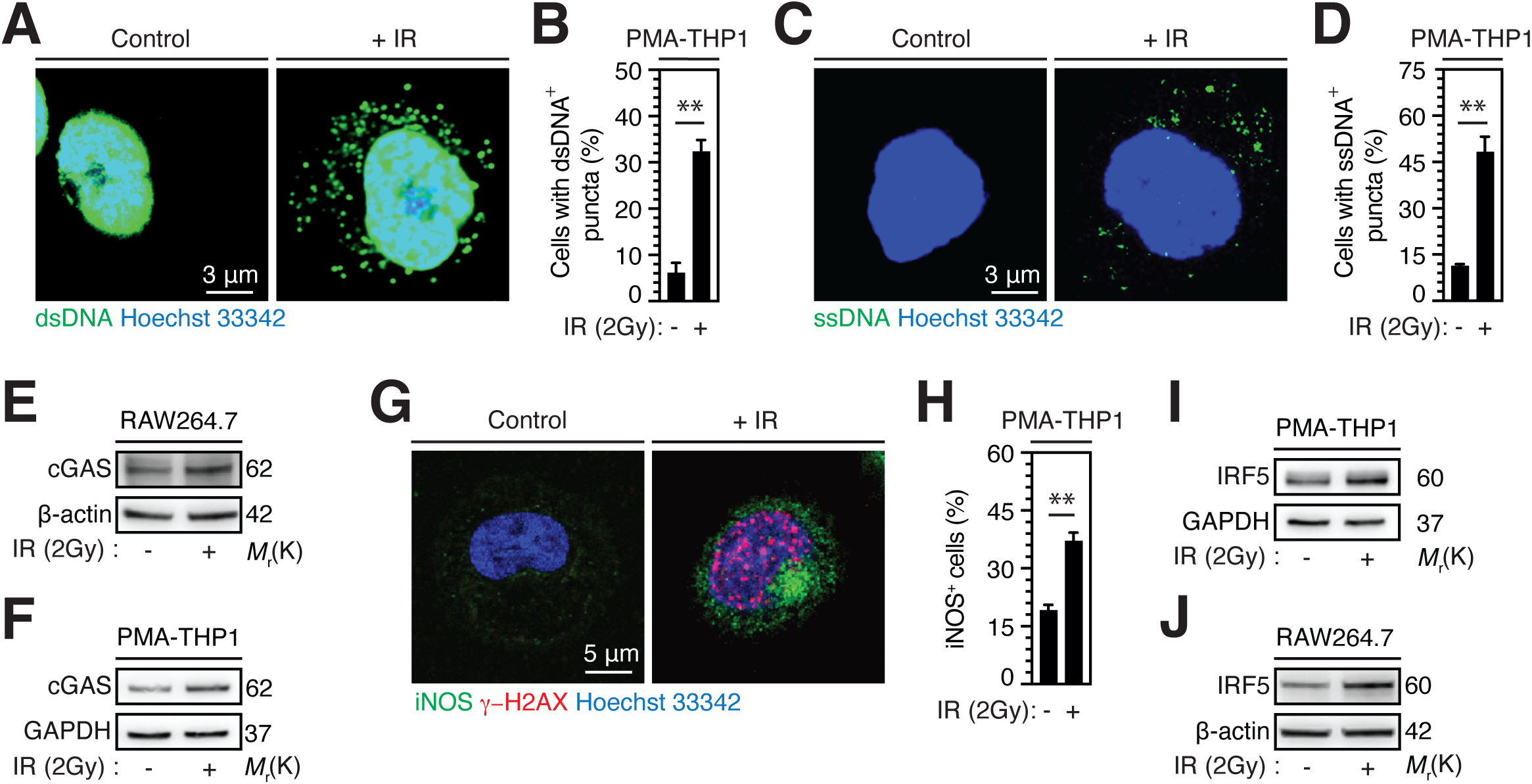
Ionizing radiation induces cytoplasmic accumulation of DNA breaks and cGAS expression during proinflammatory activation of macrophages. A, B Confocal microscopy images (A) and percentages (B) of PMA-differentiated human THP1 macrophages showing cytoplasmic double strand (ds) DNA^+^ foci 30 minutes after treatment with control or 2 Gy single-dose ionizing radiation (IR) are shown. Nuclei are stained with Hoechst 33342 (scale bar, 3 μm). C, D Confocal microscopy images (C) and percentages (D) of PMA-differentiated human THP1 macrophages showing cytoplasmic single strand (ss) DNA^+^ foci 30 minutes after treatment with control or 2 Gy single-dose IR are shown. Nuclei are stained with Hoechst 33342 (scale bar, 3 μm). E cGAS expression level detected 3 hours after the treatment of murine RAW264.7 macrophages with control or 2 Gy single-dose IR. Representative western blots are shown. β-actin is used as loading control. F cGAS expression level detected 6 hours after the treatment of PMA-differentiated human THP1 macrophages with control or 2 Gy single-dose IR. Representative western blot is shown. GAPDH is used as loading control. G, H Confocal microscopy images (G) and percentages (H) of PMA-differentiated human THP1 macrophages expressing iNOS (iNOS^+^) 24 hours after treatment with control or 2 Gy single-dose IR. Nuclei are stained with Hoechst 33342 (scale bar, 5 μm). I IRF5 expression level detected after 96 hours of treatment of PMA-differentiated human THP1 macrophages with control or 2 Gy single-dose IR. Representative western blots are shown. GAPDH is used as loading control. J IRF5 expression level detected after 6 hours of treatment of murine RAW264.7 macrophages with control or 2 Gy single-dose IR. Representative western blots are shown. β-actin is used as loading control. Images are representative from 3 independent experiments. Data are means ± S.E.M from three independent experiments. P-values (**P< 0.01) determined with two-tailed unpaired t-test (B, D, H).

### Dynamin related protein 1 (DRP1) activation triggers mitochondrial fragmentation in irradiated macrophages

Considering the central role of mitochondrial fusion/fission cycles, also referred as mitochondrial dynamics, for metabolism regulation and proinflammatory activation of macrophages [34], the shape of mitochondria in PMA-differentiated human THP1 macrophages that were irradiated with a single dose of 2 Gy was evaluated using confocal microscopy by determining the expression of the translocase of the outer mitochondrial membrane 20 (TOM20). After a single dose of 2 Gy, PMA-differentiated human THP1 macrophages exhibited a disturbed mitochondrial network and accumulated rounded, fragmented mitochondria in their cytosol (Fig. 2A), as compared to control macrophages that mainly showed an interconnected and filamentous mitochondrial network. The frequency of macrophages showing a fragmented mitochondrial network significantly increased 30 minutes after 2 Gy single-dose irradiation (Fig. 2B), as compared to control macrophages, thus revealing that mitochondrial fragmentation is rapidly induced after IR. To characterize molecular mechanisms regulating mitochondrial fragmentation in irradiated macrophages, the role of the dynamin-related protein 1 (DRP1), which is a central dynamin-related GTPase involved in the regulation of mitochondrial fission [35, 36], was evaluated during IR-mediated proinflammatory activation. Using western blots, the activating phosphorylation of DRP1 on serine 616 (DRP1S616*) was determined in 2 Gy-irradiated murine RAW264.7 macrophages (Fig. 2C) and PMA-differentiated human THP1 macrophages (Fig. 2D). The increased expression levels of DRP1S616* were observed 6 hours after the irradiation of murine RAW264.7 macrophages (Fig. 2C) or PMA-differentiated THP1 macrophages (Fig. 2D). We then appreciated the role of DRP1 activation during IR-mediated mitochondrial fragmentation and uncovered that the pharmacological DRP1 inhibitor Mdivi1 impairs the mitochondrial fragmentation detected 30 minutes after the irradiation of PMA-differentiated human THP1 macrophages with a single dose of 2 Gy (Fig. 2E-G), thus demonstrating that DRP1 activation dictates mitochondrial fragmentation in irradiated macrophages.

**Fig. 2:**
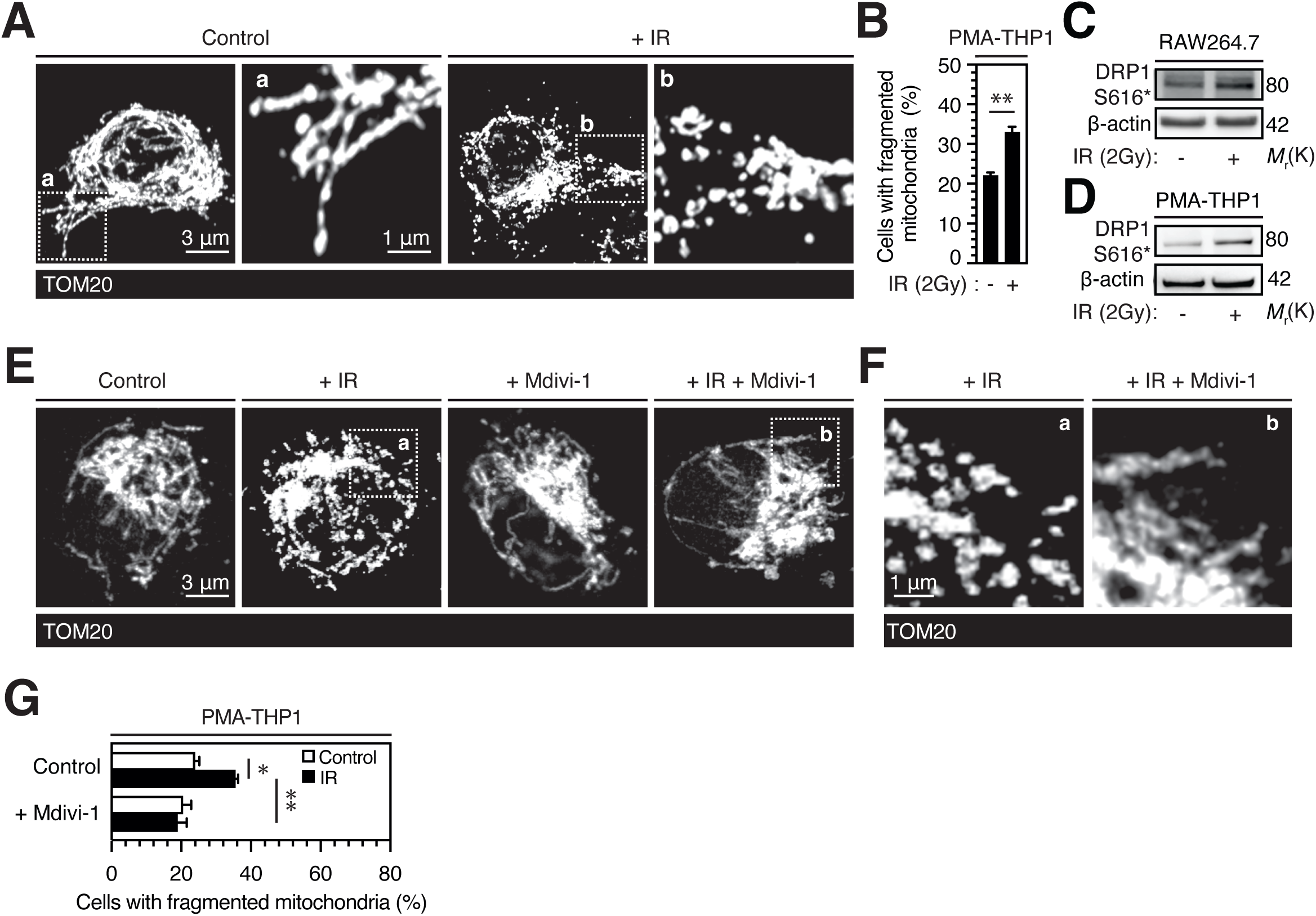
DRP1 regulates mitochondrial fragmentation during IR-mediated proinflammatory activation of macrophages. A, **B** 3D reconstructions of confocal microscopy images of the mitochondrial networks (**A**) and percentages (**B**) of PMA-differentiated human THP1 macrophages showing mitochondrial fragmentation 30 minutes after the treatment with control or 2 Gy single- dose IR are shown. Mitochondria networks are revealed by TOM 20 expression (scale bar, 3 μm). Magnifications are shown (scale bar, 1 μm). Images are representative from 3 independent experiments. **C, D** DRP1S616* expression level detected 6 hours after the treatment of murine RAW264.7 macrophages (**C**) or PMA-differentiated human THP1 macrophages (**D**) with control or 2 Gy single-dose IR. Representative western blots are shown. β-actin is used as loading control. **E** 3D reconstructions of confocal microscopy images of the mitochondrial networks of PMA- differentiated human THP1 macrophages detected 30 minutes after the treatment with 10 μM Mdivi-1 with control or 2 Gy single-dose IR are shown. Mitochondria networks are detected as shown in (**A**) (scale bar, 3 μm). Images are representative from 3 independent experiments. **F** Higher magnification details of the mitochondrial networks detected in 2 Gy-irradiated PMA- differentiated human THP1 macrophages or in PMA-differentiated human THP1 macrophages that were treated with Mdivi-1 and irradiated with a single dose of 2 Gy. a and b are magnifications of dashed regions a and b respectively shown in (**E**). Images are representative from 3 independent experiments (scale bar, 1 μm). **G** Percentages of PMA-differentiated human THP1 macrophages showing fragmented mitochondria 30 minutes after the treatment with control, 2 Gy single-dose IR, 10 μM Mdivi-1, or 2 Gy single-dose IR combined with 10 μM Mdivi-1 are shown. Data are means ± S.E.M from three independent experiments. P-values (*P< 0.05 and **P< 0.01) were determined with two-tailed unpaired t-test (**B**) or two-way ANOVA with Tukey’s multiple comparisons test (**G**).

### DRP1-mediated mitochondrial fragmentation induces IR-mediated proinflammatory macrophage activation through cGAS-dependent mechanism

Considering the ability of mitochondrial dynamics to cause cytosolic DNA accumulation directly as a result of mitochondrial DNA leakage [37] or indirectly after mitochondrial reactive oxygen species (ROS) production and genomic alteration [38], effect of the genetic inhibition of DRP1 on the increased expression level of cGAS detected 6 hours after 2 Gy single dose irradiation of PMA-differentiated human THP1 macrophages was analyzed. We observed using western blots that the depletion of DRP1 with specific small interfering RNA (siRNA) (Fig. 3A) abrogated the increase of cGAS expression level detected 6 hours after irradiation in PMA-differentiated human THP1 macrophages, thus revealing that the increase of cGAS expression level detected in irradiated macrophages depends on DRP1 activation. Finally, the role of cGAS during the IR-mediated proinflammatory macrophage activation was evaluated using confocal microscopy and western blots. PMA-differentiated human THP1 macrophages were irradiated with a single dose of 2 Gy in presence of the pharmacological inhibitor of human cGAS G140 and analyzed for iNOS, γ-H2AX and IRF5 expressions. G140 impaired the increased expression levels of iNOS (Fig. 3B, C) and IRF5 (Fig. 3D) observed 24 hours (Fig. 3B, C) and 96 hours (Fig. 3D) after IR, respectively, without affecting the accumulation of γ-H2AX^+^ foci in the nucleus of 2 Gy-irradiated PMA-differentiated human THP1 macrophages (Fig. 3B). These results indicate that cGAS acts downstream DNA damage induction in the IR-mediated proinflammatory signaling pathway that we previously described [25]. The pharmacological inhibition of murine cGAS with the use of RU521 also impaired the increased expression level of IRF5 detected 6 hours after the irradiation of murine RAW264.7 macrophages with a single-dose of 2 Gy (Fig. 3E). Accordingly, the specific depletion of cGAS using siRNA (Fig. 3F) abrogated the increased expression level of IRF5 detected 96 hours after 2Gy single-dose irradiation of PMA-differentiated human THP1 macrophages (Fig. 3G), confirming thus the central role of cGAS in the proinflammatory activation of macrophages elicited in response to IR. Altogether, these results demonstrate that IR-mediated proinflammatory macrophage activation is triggered by DRP1-dependent mitochondrial fragmentation and involves the DNA sensor cGAS.

**Fig. 3:**
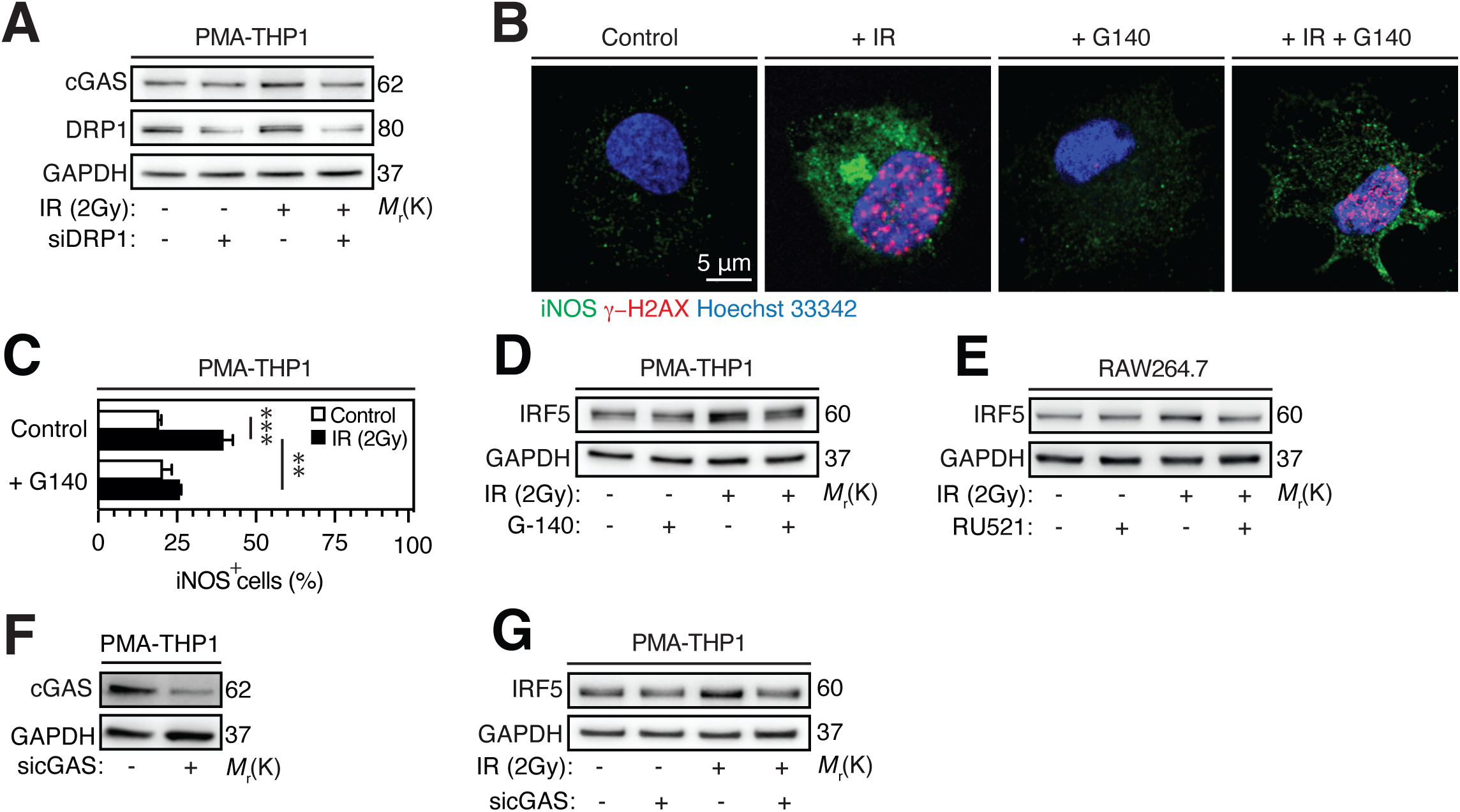
DRP1-mediated mitochondrial fragmentation dictates IR-mediated proinflammatory macrophage activation through cGAS-dependent mechanism. **A** cGAS and DRP1 expression levels after 6 hours of culture of PMA-differentiated human THP1 macrophages depleted or not for DRP1, and irradiated (or not) with a single dose of 2 Gy. Representative westerns are shown. GAPDH is used as loading control. **B, C** Confocal microscopy images (**B**) and percentages (**C**) of PMA-differentiated human THP1 macrophages showing iNOS (**B,C**) and γ-H2AX (**C**) expressions 24 hours after treatment with control, 2 Gy single-dose IR, 5 μM G140, or 2 Gy single-dose IR combined with 5 μM G140 are shown. Nuclei are stained with Hoechst 33342 (scale bar, 5 μm). Images are representative from 3 independent experiments. **D** IFR5 expression levels detected after 96 hours of culture of PMA-differentiated human THP1 macrophages that have been treated with control, 2 Gy single-dose IR, 5 μM G140, or 2 Gy single-dose IR combined with 5 μM G140. Representative western blots are shown. GAPDH is used as loading control. **E** IFR5 expression levels detected after 6 hours of culture of murine RAW264.7 macrophages that have been treated with control, 2 Gy single-dose IR, 10 μM RU521, or 2 Gy single-dose IR combined with 10 μM RU521. Representative western blots are shown. GAPDH is used as loading control. **F** cGAS expression after 96 hours of transfection with control and cGAS-specific siRNA (sicGAS). Representative western blots are shown. GAPDH is used as loading control. **G** IRF5 expression levels detected after 96 hours of culture of PMA-differentiated human THP1 macrophages depleted or not for cGAS and treated with control or 2 Gy single-dose irradiation. Representative western blots are shown. GAPDH is used as loading control. Data are means ± S.E.M from three independent experiments. P-values (**P< 0.01 and ***P< 0.001) were determined with two-way ANOVA with Tukey’s multiple comparisons test (**C**).

### Purinergic receptor P2Y2 is an endogenous repressor of proinflammatory macrophage activation

In attempt to identify combinatorial strategies to enhance IR-mediated proinflammatory activation of macrophages, we focused on the purinergic receptor P2Y2, which is membrane-anchored receptor that bind extracellular nucleotides (such as adenosine triphosphate (ATP) and uridine triphosphate (UTP)) and can regulate several macrophage functions including chemotaxis [39] and cytokine secretion [40, 41]. The impact of P2Y2 inactivation on the functional reprogramming of macrophages was then investigated. Pharmacological inhibition of P2Y2 with Kaempferol (a specific P2Y2 inhibitor) promoted classical activation of anti-inflammatory human monocyte derived macrophages (hMDMs), as revealed by the induction of known human markers of macrophage activation such as tumor necrosis factor (TNF-α), interleukin 1β (IL-1β), and interferon regulatory factor 5 (IRF5) and the reduced expression of anti-inflammatory markers such as the scavenger receptor CD163 and interleukin 10 (IL-10) (Fig. 4A). Moreover, P2Y2 inhibition with Kaempferol induced classical macrophage activation to the same extent as the positive control, interferon-γ (IFNγ) (Fig. 4B) or another purinergic receptor P2Y2 inhibitor, AR-C118925XX (AR- C) (Fig. 4C), as revealed by the increased expression of IRF5 (Fig. 4B, C). P2Y2 depletion by means of siRNAs (Fig. 4D-F) or short hairpin interfering RNA (shRNA) (Fig. 4G) increased expression levels of IRF5 (Fig. 4E-G). A diminished membrane expression of CD163 was detected on Kaempferol-treated hMDMs (Fig. 4H, I). A reduced secretion of interleukin-10 (IL-10) from PMA-differentiated human THP1 macrophages (Fig. 4J, L) or hMDMs (Fig. 4K) that were treated with Kaempferol (Fig. 4J, K) or depleted for P2Y2 (Fig. 4L) was also observed. These results indicate that P2Y2 acts as an endogenous repressor of proinflammatory macrophage activation.

**Fig. 4:**
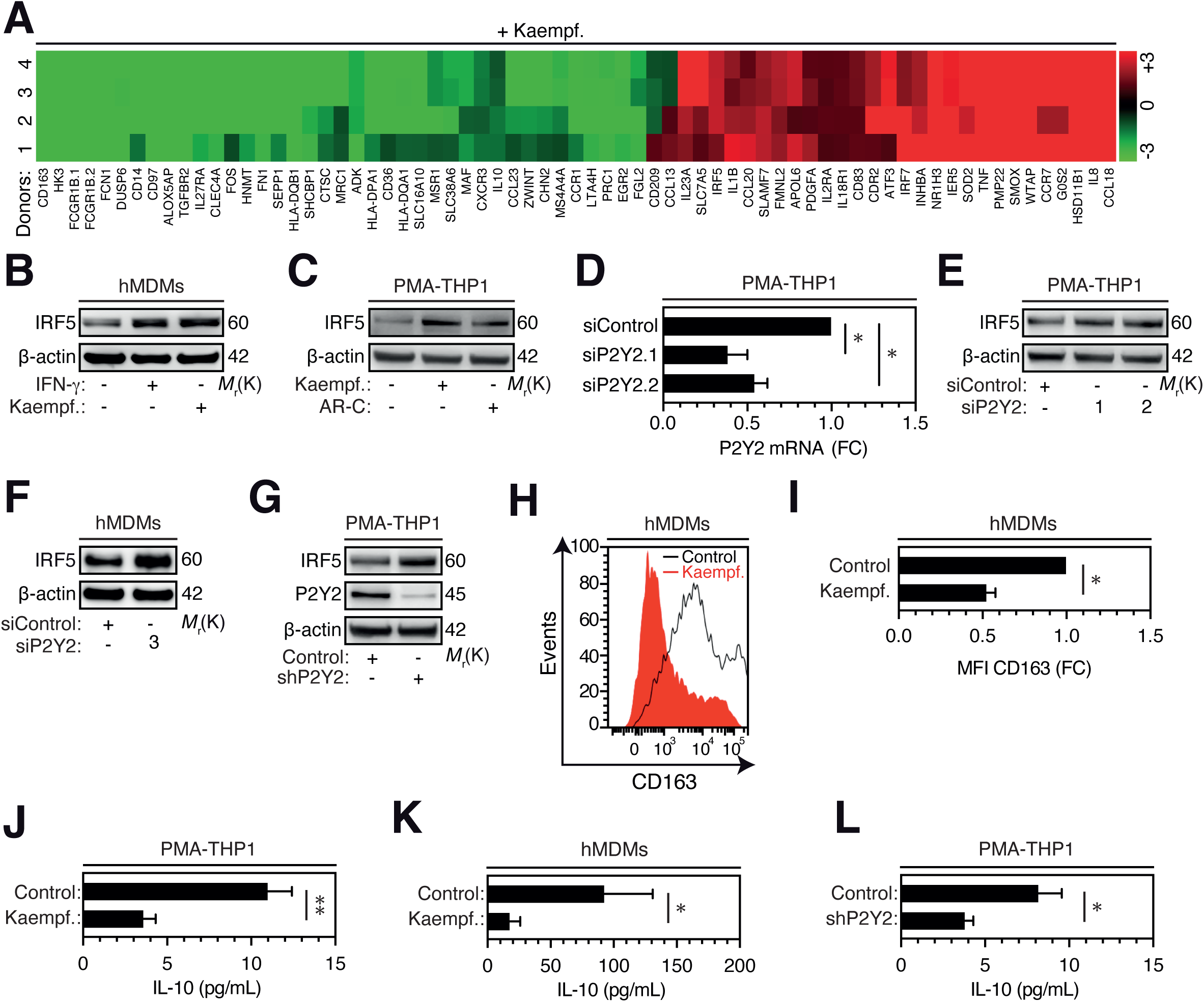
Purinergic receptor P2Y2 represses proinflammatory macrophage activation. A-L. hMDMs (**A, B, F, H, I, K**) or PMA-differentiated human THP1 macrophages (**C, D, E, G, J, L**) treated with Kaempferol (Kaempf.) (**A-C, H-K**), IFNγ (**B**), AR-C118925XX (AR-C) (**C**), transfected with siRNA (**D-F**) or expressing shRNA (**G, L**) specific for P2Y2 were analyzed for gene expression of known human polarization-specific markers (**A**), IRF5 (**B, C, E-G**), P2Y2 (**D, G**), membrane CD163 (**H, I**) and IL-10 (**J-L**) expressions using microarray assay (**A**), western blots (**B, C, E-G**), RT-PCR (**D**), flow cytometry (**H, I**) or ELISA (**J-L**). Results were obtained from at least 3 independent experiments with the exception of (**C**) with AR-C118925XX (n=2). P- values (*P< 0.05 and **P< 0.01) were determined with one sample Wilcoxon test (**D**), two-tailed Wilcoxon matched-pairs signed rank test (**I**), two-tailed unpaired t-test (**J, L**) and two-tailed Mann-Whitney test (**K**).

### P2Y2 depletion enhances IR-mediated proinflammatory activation of macrophages

Accordingly, the ability of P2Y2 inhibition to enhance IR-mediated proinflammatory macrophage activation was evaluated. The effects of P2Y2 depletion on the accumulation of DNA breaks and fragmented mitochondria in the cytosol of 2Gy-irradiated PMA-differentiated human THP1 macrophages were first studied. We observed that P2Y2 depletion is sufficient to induce the accumulation of both dsDNA and ssDNA fragments in the cytosol of P2Y2-depleted macrophages (Fig. 5A-D). Despite the fact that the frequency of macrophages showing ssDNA-positive puncta did not vary between macrophages depleted for P2Y2 that were irradiated or not with a single dose of 2 Gy (Fig. 5D), the frequency of macrophages showing dsDNA-positive puncta was significantly higher in 2Gy-irradiated macrophages that were depleted for P2Y2 than in 2Gy-irradiated control macrophages (Fig. 5B). As shown in Figures 1 and 2, the cytosolic accumulation of DNA breaks positively correlates with the presence of fragmented mitochondria detected in 2Gy-irradiated macrophages, in P2Y2-depleted macrophages and in 2Gy-irradiated macrophages that were depleted for P2Y2 (Fig. 5E, F). Interestingly, we also revealed that P2Y2 depletion or 2Gy single-dose irradiation increased the frequency of macrophages expressing iNOS (Fig. 5G) and showed that P2Y2 depletion significantly increased the ability of 2Gy-irradiated PMA-differentiated human THP1 macrophages to express iNOS (Fig. 5G, H), as compared to control or 2Gy-irradiated PMA-differentiated human THP1 macrophages. Altogether, these results demonstrate that P2Y2 depletion enhances the ability of irradiated macrophages to undergo a proinflammatory activation.

**Fig. 5:**
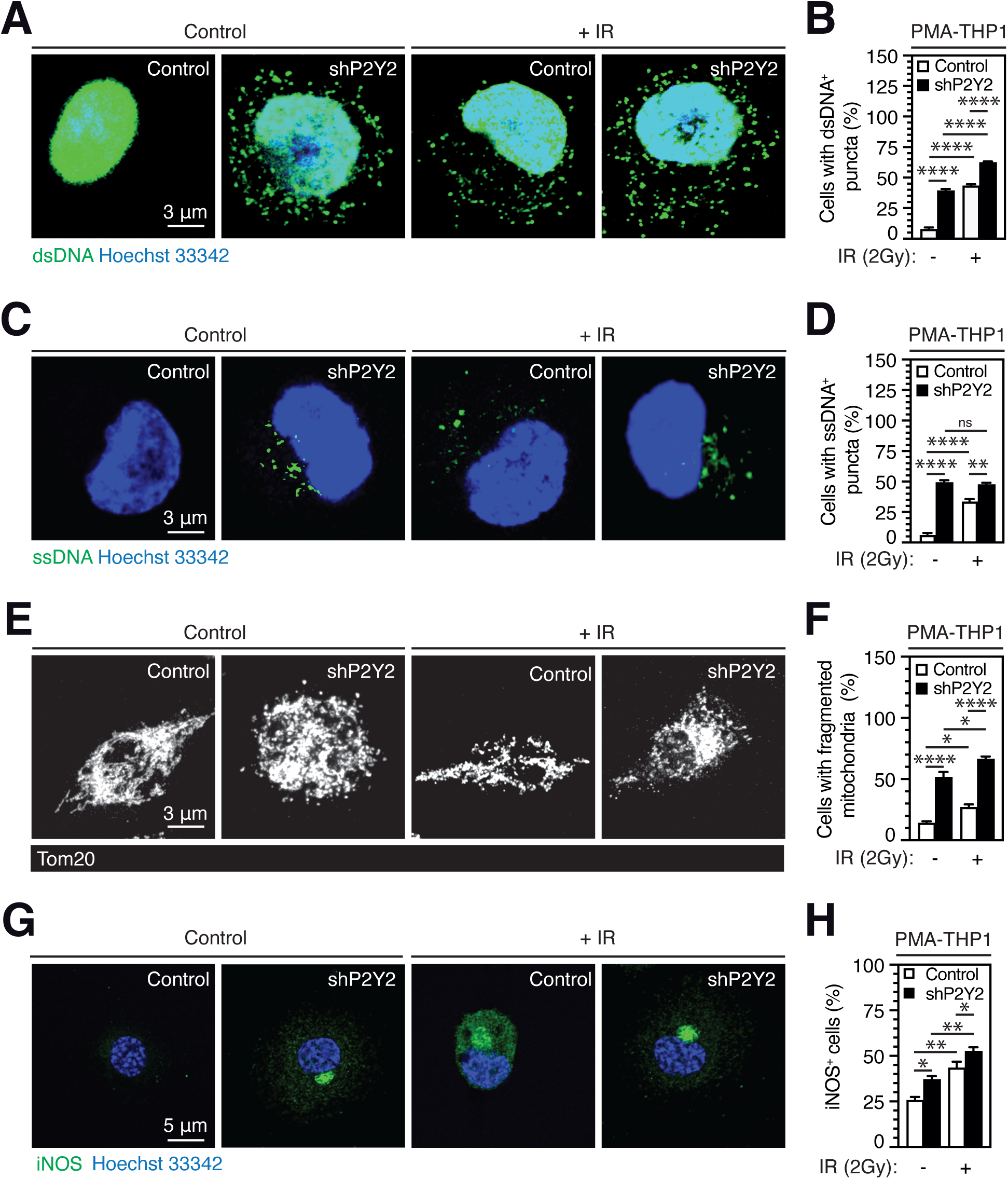
P2Y2 depletion enhances IR-mediated proinflammatory macrophage activation. A-F. Confocal microscopy images (**A, C, E**) and percentages (**B, D, F**) of PMA-differentiated, control or P2Y2-depleted human THP1 macrophages showing cytoplasmic dsDNA^+^ foci (**A, B**), ssDNA^+^ foci (**C, D**) or fragmented mitochondria (**E, F**) 30 minutes after treatment with control or 2 Gy single-dose ionizing radiation (IR) are shown. Nuclei are stained with Hoechst 33342 (scale bar, 3 μm). **G, H** Confocal microscopy images (**G**) and percentages (**H**) of PMA-differentiated, control or P2Y2-depleted human THP1 macrophages expressing (or not) iNOS (iNOS^+^) after 24-hour treatment with control or 2 Gy single-dose IR are shown. Nuclei are stained with Hoechst 33342 (scale bar, 5 μm). Data are means ± S.E.M from three independent experiments. P-values (*P< 0.05, **P< 0.01 and ****P< 0.0001) were determined with two-way ANOVA with Tukey’s multiple comparisons test (**B, D, F, H**).

## Discussion

Despite the fact that radiotherapy is a cornerstone of cancer treatment, the molecular and cellular bases of radiotherapy efficacy are still poorly understood and the characterization of antitumor immune-modulatory activities elicited by radiotherapy is under intensive investigations. We previously demonstrated that IR promotes the proinflammatory activation of macrophages in a cell-autonomous manner through the increased expression of NOX2 and ROS production [25]. Oxidative stress causes DNA damages in the nucleus of irradiated macrophages and subsequently leads to the proinflammatory activation of irradiated macrophages through ATM kinase activation [25]. We also reported that the inhibition of this signaling pathway impairs the proinflammatory activation of macrophages and predicts tumor response to hypofractionated radiotherapy in locally advanced rectal cancer [25], thus confirming that the reprogramming of anti-inflammatory TAMs into proinflammatory TAMs is associated with the efficacy of radiotherapy. In this context and with the ambition to identify novel cellular targets for the improvement of radiotherapy efficacy, we further explored molecular mechanisms regulating the proinflammatory activation of macrophages in response to IR.

In this paper, we show that IR rapidly triggered the cytosolic accumulation of single and double stranded DNA breaks, and the stabilization of DNA sensor cGAS in irradiated macrophages, thus suggesting that cGAS may sense IR-induced self-DNA breaks that accumulated in the cytosol of irradiated macrophages and potentially participate to the proinflammatory activation of irradiated macrophages. To determine the causal relationship between the cytosolic accumulation of self-DNA and proinflammatory macrophage activation, the role of cGAS in this process was studied. Our results revealed that in response to IR, cGAS induces the increased expression of iNOS and IRF5 and thus, favors the proinflammatory activation of macrophages. Our results are consistent with previous publication revealing that cytosolic DNA sensing by cGAS governs macrophage proinflammatory activation in myocardial ischemic injury [41]. Interestingly, cGAS inhibition did not affect the accumulation of γ-H2AX-containing nuclear foci, thus indicating that cGAS may act after the induction of DNA damage and the ATM-dependent DNA damage response that we previously reported during the IR-mediated proinflammatory macrophage activation [25]. Considering that the recognition of cytosolic DNA [29] and the increased abundance of cGAS mRNA [42] can both lead the increased expression level of cGAS, further investigations are needed to define the molecular mechanisms involved in the upregulation of cGAS expression levels detected in macrophages in response to IR.

We also noticed that fragmented mitochondria accumulated in macrophages upon irradiation, suggesting that macrophage mitochondrial dynamics can be affected by IR and may potentially contribute to the proinflammatory activation of macrophages. The increased activating phosphorylation of the major pro-fission protein DRP1 (DRP1S616*) was detected rapidly after the irradiation of macrophages. Moreover, the pharmacological inhibition and the depletion of DRP1 respectively reduced mitochondrial fragmentation and cGAS expression levels in irradiated macrophages, thus highlighting that DRP1 activation is required for IR-mediated mitochondrial fragmentation and dictates cGAS expression level during proinflammatory activation of irradiated macrophages. Our results are in line with the ability of DRP1 to promote mitochondrial fragmentation and regulate the secretion of IL-12, IL-6 and IL-1β during LPS-induced proinflammatory macrophage activation [43]. Considering the ability of protein kinase C δ [44], cyclin-dependent kinases 1 (CDK1) and 5 (CDK5) [44, 45], calcium–calmodulin (CaM)-dependent protein kinase II [46], extracellular signal regulated kinase ERK2 [47] and PTEN-induced putative kinase 1 (PINK1) [48] to induce DRP1S616* phosphorylation, their contribution to IR-mediated proinflammatory macrophage activation remain to be addressed. Moreover, molecular mechanisms linking DRP1-dependent mitochondrial fragmentation and the increased expression levels of cGAS expression remain to be defined. Our results also reveal that cGAS supports IR-mediated proinflammatory macrophage activation, corroborating recent results showing that the cGAS regulates mTORC1 pathway and controls proinflammatory macrophage activation during sepsis-induced lung injury [49] or that the activation of STING with the agonist DXMAA promotes a shift from anti-inflammatory to proinflammatory phenotype of TAMs [50]. Accordingly, a particular attention should be paid to the messenger 2′,3′-cGAMP, the kinase STING, other transcription factors such as IRF3 and NF-κB and the type 1 interferon secretion (particularly IFN-β) that are known to act downstream to cGAS [29, 51]. Our findings thus provide insights into the role of cGAS during radiotherapy and demonstrate that IR triggers a proinflammatory signaling pathway in macrophages involving DRP1S616*, mitochondrial dynamics, cytosolic DNA fragments, cGAS and IRF5. The crosstalk between this signaling pathway and the molecular cascade (involving NOX2, ROS production, ATMS1981* and IRF5) that we previously identified [25], need to be addressed. Considering that ATM kinase was shown to restrain spontaneous type I Interferon [52], its contribution to IR-mediated proinflammatory macrophage activation need further investigations.

Our results also provide strong evidence that purinergic receptor P2Y2-mediated signaling pathways play a central role in the repression of proinflammatory macrophage activation. Pharmacological inhibition (with Kaempferol or AR-C119825XX) or depletion of P2Y2 promotes the proinflammatory activation of PMA-differentiated human THP1 macrophages and murine RAW264.7 macrophages. More importantly, the inactivation of P2Y2 reprogrammed anti-inflammatory hMDMs to proinflammatory hMDMs, thus revealing that P2Y2-mediated signaling pathways negatively regulate the switch from anti-inflammatory macrophages to proinflammatory macrophages. Our results are consistent with the effects of pharmacological and macrophage-specific genetic inhibition of the β-catenin [53], the PI3-kinase γ [21, 54] or the transcription factor c-MYC [55], which are critical for the anti-inflammatory reprogramming of macrophages. Considering that purinergic receptor P2Y2 has been involved in regulation of the biological activities of β-catenin [56], c-MYC [57] and the PI3K/AKT-dependent signaling pathway [58], further investigations should analyze effects of P2Y2 inactivation on the transcriptional activation of FOS-like antigen 2 (FOSL2) and NF−κB, and on the repression of the AT-rich interaction domain 5A (ARDI5A) or C/EBPβ activation, which are critical for macrophage functional reprogramming elicited by the inhibition of β-catenin, the PI3-kinase γ or c-MYC [21, 53, 55].

We and others proposed that the reprogramming of TAMs with IR alone or in combination with other modalities of cancer treatments (such as PARP inhibitors [59] or functionalized nanoparticles [60]) might enhance the efficacy of radiotherapy [15]. Accordingly, the identification of novel mechanisms regulating macrophage functions could help to identify novel components whose therapeutic modulations should improve the proinflammatory activation of macrophages and the efficacy of radiotherapy. Here, we identify P2Y2 as an endogenous repressor of proinflammatory macrophage activation and demonstrated that the combination of IR with P2Y2 inactivation enhanced the ability of macrophages to accumulate dsDNA breaks and fragmented mitochondria in their cytosol and to be proinflammatory activated in response to IR. Our results suggest that combinatorial strategies involving IR and P2Y2 inactivation should stimulate the proinflammatory signaling pathway (involving DRP1S616*, mitochondrial dynamics, cytosolic DNA fragments, cGAS and IRF5) that we described above. Further molecular characterizations are needed to precise the molecular cascade involved in this process. Beside these functional considerations, the identification of innovative therapeutic strategies allowing the specific inhibition of P2Y2 in TAMs might help to improve patient’s response to radiotherapy and have the potential to synergize with other cancer treatments such as immunotherapies.

## Materials and methods

### Cells and Reagents

The human monocytic THP1 cell line and murine RAW264.7 macrophages were obtained from ATCC and respectively cultured in RMPI-1640-Glutamax medium (Life Technologies, USA) and in DMEM-Glutamax medium (Life Technologies, Carlsbad, USA) supplemented with 10% heat-inactivated fetal bovine serum (HI FBS) (Hycultec GmbH, Beutslsbach, Germany) and 100 IU/ml penicillin–streptomycin (Life Technologies). Cells were maintained at 37 °C under 5% CO2 humidified atmosphere. Human THP1 cells were differentiated into THP1 macrophages with 320 nM of PMA (#tlrl-PMA, Invivogen, USA) during 24 hours and cultured for additional 24 hours before use. Human monocyte-derived macrophages (hMDMs) were obtained and differentiated as previously described [61]. Kaempferol (#K0133) and AR-C118925XX (#4890) were obtained from Sigma-Aldrich (L’Isle-d’Abeau, France) and Tocris (Bristol, UK), respectively. Recombinant Human IFN-*γ* (IFN-*γ*, #285-IF/CF) was from R&D Systems (Minneapolis, MN, USA). Mdivi-1 (#S7162) was from Selleckchem (Houston, TX, USA). G140 (#inh-g140) and RU521 (#inh-ru- 521) were from Invivogen (Toulouse, France).

### Antibodies

Antibodies used for immunofluorescence were anti-phospho-H2AX (Ser139) (#05-636) antibody from EMD Millipore (Billerica, MA, USA), anti-iNOS (#ab3523) and anti-dsDNA (#ab27156) antibodies from Abcam (Cambridge, UK), anti-Tom20 (F-10) (#sc-17764) antibody from Santa Cruz Biotechnology (Oregon, USA) and anti-DNA, single stranded specific, clone F7-26 (MAB3299) antibody from Merck (Darmstadt, DE). For western blots, anti-cGAS (D1D3G) (#15103), anti-cGAS (D3O8O) (#31659), anti-Phospho-DRP1 (Ser616) (#3455) antibodies were from Cell Signaling Technology. Anti-P2Y2 antibody (#APR-010) was from Alomone laboratories. Anti-IRF5 (#ab21689) and anti-beta Actin HRP [AC-15] (#ab49900) antibodies were from Abcam, and anti-GAPDH (#MAB374) antibody was from BD bioscience. Anti-CD163 Alexa Fluor 647 (#562669) was from BD Pharmingen.

### Irradiation

As described above, human THP1 cells were differentiated into macrophages with 320 nM of PMA (Invivogen, #tlrl-PMA) in 12-well plates (10^6^ cells/well) or in 24-well plates (5x10^5^ cells/well). Murine RAW264.7 macrophages were seeded in 24-well plates (5x10^4^ cells/well) 24 hours before treatments. Macrophages were then irradiated with a single dose of 2 Gy in presence or in absence of indicated pharmacological inhibitors or after indicated knockdowns with X-ray irradiator (1 Gy/min, X-RAD 320, Precision X-Ray). Macrophages were harvested at 30 minutes, 1 hour, 6 hours, 24 hours or 96 hours after irradiation for western blot and/or immunofluorescence analyses.

### siRNA and shRNA for gene silencing

PMA-differentiated human THP1 macrophages were transfected using specific siRNAs for human P2Y2 (siP2Y2.1 et siP2Y2.2) or control siRNAs (siControl) from Sigma-Aldrich, SMARTpool siGENOME Human MB21D1 (115004) siRNAs (M-015607-01-0005) for cGAS, ON-TARGET plus Human DNM1L (10059) siRNAs for DRP1, siGENOME Non-Targeting siRNA Pool #2 (D- 001206-14-05) or ON-TARGET plus Non-targeting Pool (D001810-10-05) as controls from Dharmacon (Lafayette, CO, USA). Lipofectamine RNAi max (#13778150, Life Technologies, Illkrich, France) was used for transfection, according to the manufacturer’s instructions. Briefly, Lipofectamine RNAi max and siRNA were mixed in Opti-MEM medium (Thermo Fisher Scientific) without serum and were incubated for 5 minutes at room temperature. The transfection mix was then added to PMA-differentiated human THP1 macrophages for 48 hours at 37 °C. After 48 hours, the medium was replaced by fresh medium supplemented with 10% FBS and transfected macrophages were irradiated. For the transfection of hMDMs, smart pools of siGenome non-targeting control and P2Y2 specific siRNAs (siP2Y2.3) were obtained from Dharmacon and used as previously described [25, 62, 63]. Control and P2Y2-depleted (shP2Y2) THP1 monocytes were used as previously described [62]. Sequences of human siRNAs, siGENOME SMARTpool and shRNA used in this study are shown in Supplementary Table 1.

### Immunofluorescence and flow cytometry

PMA-differentiated human THP1 macrophages were cultured on coverslips and fixed after treatment with 4% (w/v) paraformaldehyde (Sigma-Aldrich, #1.000496.500) in Dubelco’s phosphate buffer saline (DPBS) (Gibco, #14190-094) for 5 minutes. Macrophages were permeabilized with 0.3% Triton X-100 (Sigma-Aldrich, #X100) in DPBS for 5 minutes at room temperature, washed in DPBS and blocked with 10% HI FBS in DPBS for 1 hour. Macrophages were then incubated with primary antibodies in 10% HI FBS in DPBS for 2 hours, washed with DPBS and incubated with the secondary antibodies Alexa Fluor 488 (Invitrogen, #A11034) or Alexa Fluor-546 (Invitrogen, #A11030) and Hoechst 33342 (Invitrogen, #1874027) in DPBS containing 10% HI FBS for 30 minutes at room temperature. Coverslips were mounted with Fluoromount-G (SouthernBiotech, Birmingham, AL, USA) on microscope slides (Thermo Fisher Scientific) and examined with Leica TCS SPE confocal microscope (Leica Microsystems, Nanterre, France) using a X63 oil objective, as previously described [63]. After indicated treatments, hMDMs (10^6^ cells/mL) were harvested in RPMI complete medium, washed with DPBS, saturated at 4°C for 20 minutes in DPBS containing 10% HI FBS and incubated with anti- CD163 Alexa Fluor 647 (BD Pharmingen, #562669) antibody for 1 hour and 30 minutes. Membrane expression of CD163 was then analyzed using LSRFortessa flow cytometer (BD Biosciences).

### Western blot analysis

Western blots were performed as previously published [25, 61, 62, 63]. Total cellular proteins were extracted from treated or transfected PMA-differentiated human THP1 macrophages, murine RAW264.7 macrophages or hMDMs in lysis buffer containing 1 M Tris (pH = 7.4), 5 M NaCl, 0.1% CHAPS, and protease and phosphatase inhibitors (Roche, Basel, Switzerland). Protein extracts (3-15 μg) were separated on 10%, 12% or 4-12% NuPAGE gels (Invitrogen, Illkrich, France) and transferred onto 0.2 μm pore nitrocellulose membranes (Bio-Rad, Marnes-la-coquette, France) at room temperature. After saturation with in Tris-buffered saline containing 0.1% Tween 20 (TBS-T) and 5% bovine serum albumin, membranes were incubated with indicated primary antibodies overnight at 4 °C and with horseradish peroxidase-conjugated anti-rabbit or anti-mouse IgG (SouthernBiotech, Birmingham, AL, USA) for 2 hours at room temperature. Nitrocellulose membranes were then washed with TBS-T and analyzed with enhanced ECL detection system (GE Healthcare).

### Microarray assay, data processing and analysis

Human MDMs were treated with 100 μM Kaempferol during 72 hours. Then, mRNAs were isolated using RNeasy kit (#74104, Quiagen) and gene expression analyses were performed with Agilent® SurePrint G3 Human GE 8x60K Microarray (Agilent Technologies, AMADID 39494) with the following single-color design. RNAs were labeled with Cy3 using the one-color Agilent labeling kit (Low Input Quick Amp Labeling Kit 5190-2306) adapted for small amount of total RNA (100 ng total RNA per reaction). Hybridization was then performed on microarray using 800 ng of linearly amplified cRNA labeled, following the manufacturer protocol (Agilent SureHyb Chamber; 800 ng of labeled extract; duration of hybridization of 17 hours; 40 µL per array; Temperature of 65 °C). After washing in acetonitrile, slides were scanned by using an Agilent G2565 C DNA microarray scanner with defaults parameters (100° PMT, 3 µm resolution, at 20°C in free ozone concentration environment). Microarray images were analyzed by means of Feature Extraction software version (10.7.3.1) from Agilent technologies. Defaults settings were used. Raw data files from Feature Extraction were imported into R with LIMMA (Smyth, 2004, Statistical applications in Genetics and molecular biology, vol 3, N°1, article 3), an R package from the Bioconductor project, and processed as follow: gMedianSignal data were imported, controls probes were systematically removed, and flagged probes (gIsSaturated, gIsFeatpopnOL, gIsFeatNonUnifOL) were set to NA. Inter-array normalization was performed by quantile normalization. To get a single value for each transcript, taking the median of each replicated probes summarized data. Missing values were inferred using KNN algorithm from the package ‘impute’ from R bioconductor. Normalized data were then analyzed. To assess differentially expressed genes between two groups, we started by fitting a linear model to the data. Then, we used an empirical Bayes method to moderate the standard errors of the estimated log-fold changes. The top-ranked genes were selected with the following criteria: an absolute fold-change > 2 and an adjusted p-value (FDR) < 0.05.

### Detection of IL10 release

Supernatants harvested from hMDMs or PMA-differentiated human THP1 macrophages that were treated (or not) with 100 μM Kaempferol at indicated times, or from PMA-THP1 that were depleted (or not) for P2Y2 were analyzed using ELISA for IL-10 (BD Biosciences, # 550613), according to the manufacturers’ instructions.

### Quantitative RT-PCR

After treatment, total cell RNA was extracted using NucleoSpin RNA Plus XS, Micro kit for RNA purification with DNA removal column (#740990.250, Macherey-Nagel) according to the manufacturer’s instructions. RNA was submitted to reverse transcription and PCR amplification. Quantifications were performed by real-time PCR on a Light Cycler instrument (Roche Diagnostics, Meylan, France) or using CFX Maestro (BioRad). The amplification of human P2Y2 (Catalog #4331182 Assay ID Hs01856611_s1, Thermo Fisher Scientific) and human β-actin (Catalog #4331182 Assay ID Hs01060665_g1, Thermo Fisher Scientific) was obtained after 5 minutes of reverse transcription, 5 minutes of denaturation and 45 to 50 cycles with the following steps (95°C during 5 seconds, 60°C during 15 seconds and 72°C during 15 seconds or 95°C for 15 seconds and 58°C for 30 seconds to 1 minute). Cq results were normalized by β-actin mRNA expression levels and reported to control condition. Data are presented as fold changes (FC) and were calculated with relative quantification of ΔCT obtained from quantitative RT-PCR.

## Statistical analysis

Statistical analysis was performed using GraphPad Prism 8.1 (GraphPad). Statistical tests and calculated P-values (*P< 0.05, **P< 0.01, ***P< 0.001, and ****P< 0.0001) are indicated in each figure and corresponding figure legend.

## Acknowledgments

We gratefully acknowledge the Imaging and Cytometry Platform (Y. Lecluse, S. Salome- Desnoulez and Tudor Manoliu), IMS 3655 CNRS / US 23 INSERM (Gustave Roussy, Villejuif, France) and Bioinformatics core facility (Guillaume Meurice) from Gustave Roussy research platforms for their technical support.

## Author contribution statement

Jean-Luc Perfettini provided financial support. Jean-Luc Perfettini designed and conducted the study. Ali Mostefa-Kara, Aurélia Deva Nathan, Audrey Paoletti, Anais Chapel, Emie Gutierrez- Mateyron, Thanh Trang Cao-Pham and Awatef Allouch performed experiments. Ali Mostefa-Kara, Aurélia Deva Nathan, Audrey Paoletti, Anais Chapel, Emie Gutierrez-Mateyron, Thanh Trang Cao-Pham, Yara Bou Ghosn, Awatef Allouch and Jean-Luc Perfettini analyzed the results. Ali Mostefa-Kara, Aurélia Deva Nathan, Audrey Paoletti and Anais Chapel performed statistical analysis. Ali Mostefa-Kara, Aurélia Deva Nathan, Audrey Paoletti, Anais Chapel, Emie Gutierrez- Mateyron and Awatef Allouch assembled the figures. Ali Mostefa-Kara, Aurélia Deva Nathan, Awatef Allouch and Jean-Luc Perfettini wrote the initial draft. Audrey Paoletti, Anais Chapel, Emie Gutierrez-Mateyron, Thanh Trang Cao-Pham and Yara Bou Ghosn provided advice and edited the initial draft. All authors read and approved the final version.

## Supplementary information

**Supplementary Figure S1.**
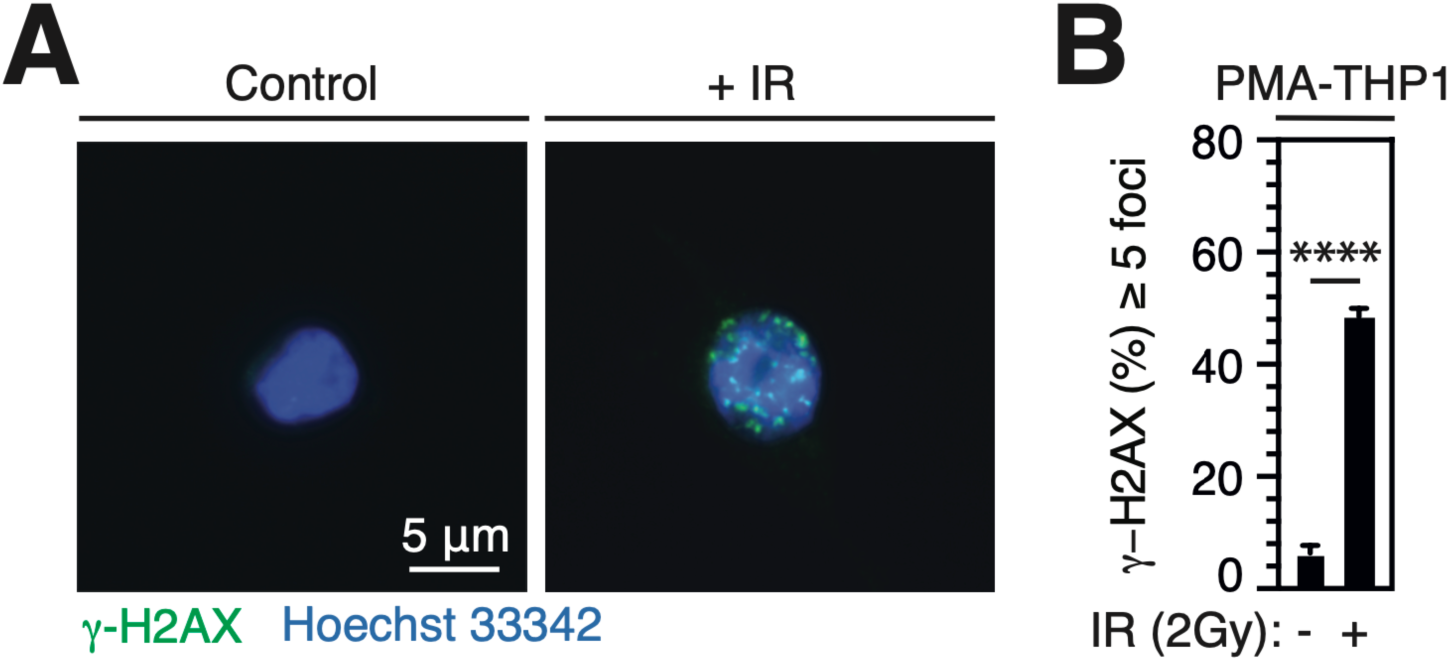
: **PMA-THP1 macrophages exhibit γ-H2AX^+^ foci after 2-Gy irradiation. A, B** Fluorescence microscopy images (**A**) and percentages (**B**) of PMA- differentiated human THP1 macrophages showing γ-H2AX^+^ nuclear foci 1 hour after treatment with control or 2 Gy single-dose IR are shown (scale bar, 5 μm). Nuclei are stained with Hoechst 33342. Data are means ± S.E.M from three independent experiments. P-values (****P< 0.0001) were determined with two-tailed unpaired t test (**B**).

**Supplementary Table S1.**
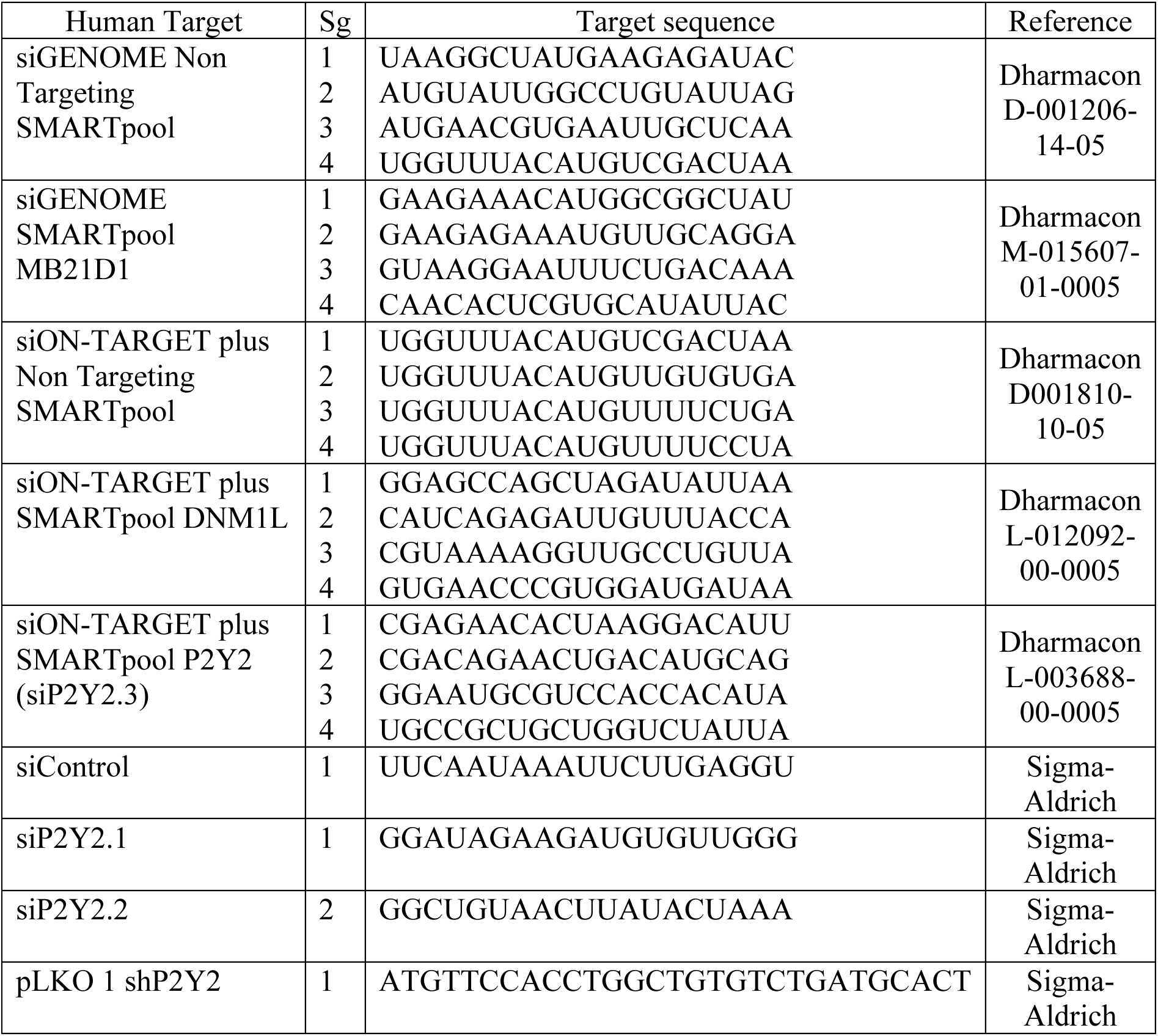
Human siRNA, siGENOME SMARTpool, siON-TARGETplus SMARTpool and shRNA used in this study.

## References

1. Baumann M, Krause M, Overgaard J, Debus J, Bentzen SM, Daartz J, et al. Radiation oncology in the era of precision medicine. Nat Rev Cancer 2016; 16: 234–249.

2. Martins I, Raza SQ, Voisin L, Dakhli H, Law F, De Jong D, et al. Entosis: The emerging face of non-cell-autonomous type IV programmed death. Biomed J 2017; 40: 133–140.

3. Verheij M, Bartelink H. Radiation-induced apoptosis. Cell Tissue Res 2000; 301: 133–142.

4. Daido S, Yamamoto A, Fujiwara K, Sawaya R, Kondo S, Kondo Y. Inhibition of the DNA- dependent protein kinase catalytic subunit radiosensitizes malignant glioma cells by inducing autophagy. Cancer Res 2005; 65: 4368–4375.

5. Huang Z, Epperly M, Watkins SC, Greenberger JS, Kagan VE, Bayir H. Necrostatin-1 rescues mice from lethal irradiation. Biochim Biophys Acta 2016; 1862: 850–856.

6. Adjemian S, Oltean T, Martens S, Wiernicki B, Goossens V, Vanden Berghe T, et al. Ionizing radiation results in a mixture of cellular outcomes including mitotic catastrophe, senescence, methuosis, and iron-dependent cell death. Cell Death Dis 2020; 11: 1003.

7. Martins I, Raza SQ, Voisin L, Dakhli H, Allouch A, Law F, et al. Anticancer chemotherapy and radiotherapy trigger both non-cell-autonomous and cell-autonomous death. Cell Death Dis 2018; 9: 716.

8. Ye LF, Chaudhary KR, Zandkarimi F, Harken AD, Kinslow CJ, Upadhyayula PS, et al. Radiation-Induced Lipid Peroxidation Triggers Ferroptosis and Synergizes with Ferroptosis Inducers. ACS Chem Biol 2020; 15: 469–484.

9. Ianzini F, Mackey MA. Spontaneous premature chromosome condensation and mitotic catastrophe following irradiation of HeLa S3 cells. Int J Radiat Biol 1997; 72: 409–421.

10. Jones KR, Elmore LW, Jackson-Cook C, Demasters G, Povirk LF, Holt SE, et al. p53- Dependent accelerated senescence induced by ionizing radiation in breast tumour cells. Int J Radiat Biol 2005; 81: 445–458.

11. Martins I, Tesniere A, Kepp O, Michaud M, Schlemmer F, Senovilla L, et al. Chemotherapy induces ATP release from tumor cells. Cell Cycle 2009; 8: 3723–3728.

12. Scaffidi P, Misteli T, Bianchi ME. Release of chromatin protein HMGB1 by necrotic cells triggers inflammation. Nature 2002; 418: 191–195.

13. Deng L, Liang H, Xu M, Yang X, Burnette B, Arina A, et al. STING-Dependent Cytosolic DNA Sensing Promotes Radiation-Induced Type I Interferon-Dependent Antitumor Immunity in Immunogenic Tumors. Immunity 2014; 41: 843–852.

14. Burnette BC, Liang H, Lee Y, Chlewicki L, Khodarev NN, Weichselbaum RR, et al. The efficacy of radiotherapy relies upon induction of type i interferon-dependent innate and adaptive immunity. Cancer Res 2011; 71: 2488–2496.

15. Wu Q, Allouch A, Martins I, Brenner C, Modjtahedi N, Deutsch E, et al. Modulating Both Tumor Cell Death and Innate Immunity Is Essential for Improving Radiation Therapy Effectiveness. Front Immunol 2017; 8: 613.

16. Barker HE, Paget JT, Khan AA, Harrington KJ. The tumour microenvironment after radiotherapy: mechanisms of resistance and recurrence. Nat Rev Cancer 2015; 15: 409–425.

17. Biswas SK, Mantovani A. Macrophage plasticity and interaction with lymphocyte subsets: cancer as a paradigm. Nat Immunol 2010; 11: 889–896.

18. Feng M, Jiang W, Kim BYS, Zhang CC, Fu YX, Weissman IL. Phagocytosis checkpoints as new targets for cancer immunotherapy. Nat Rev Cancer 2019; 19: 568–586.

19. Mantovani A, Marchesi F, Malesci A, Laghi L, Allavena P. Tumour-associated macrophages as treatment targets in oncology. Nat Rev Clin Oncol 2017; 14: 399–416.

20. Advani R, Flinn I, Popplewell L, Forero A, Bartlett NL, Ghosh N, et al. CD47 Blockade by Hu5F9-G4 and Rituximab in Non-Hodgkin’s Lymphoma. N Engl J Med 2018; 379: 1711–1721.

21. Kaneda MM, Cappello P, Nguyen AV, Ralainirina N, Hardamon CR, Foubert P, et al. Macrophage PI3Kgamma Drives Pancreatic Ductal Adenocarcinoma Progression. Cancer Discov 2016; 6: 870–885.

22. Zhang P, Miska J, Lee-Chang C, Rashidi A, Panek WK, An S, et al. Therapeutic targeting of tumor-associated myeloid cells synergizes with radiation therapy for glioblastoma. Proc Natl Acad Sci U S A 2019; 116: 23714–23723.

23. Klug F, Prakash H, Huber PE, Seibel T, Bender N, Halama N, et al. Low-dose irradiation programs macrophage differentiation to an iNOS(+)/M1 phenotype that orchestrates effective T cell immunotherapy. Cancer Cell 2013; 24: 589–602.

24. Wu Q, Allouch A, Martins I, Modjtahedi N, Deutsch E, Perfettini JL. Macrophage biology plays a central role during ionizing radiation-elicited tumor response. Biomed J 2017, 40: 200–211.

25. Wu Q, Allouch A, Paoletti A, Leteur C, Mirjolet C, Martins I, et al. NOX2-dependent ATM kinase activation dictates pro-inflammatory macrophage phenotype and improves effectiveness to radiation therapy. Cell Death Differ 2017; 24: 1632–1644.

26. Huang RX, Zhou PK. DNA damage response signaling pathways and targets for radiotherapy sensitization in cancer. Signal Transduct Target Ther 2020; 5: 60.

27. Yamazaki T, Kirchmair A, Sato A, Buque A, Rybstein M, Petroni G, et al. Mitochondrial DNA drives abscopal responses to radiation that are inhibited by autophagy. Nat Immunol 2020; 21: 1160–1171.

28. Oliveira M, Rodrigues DR, Guillory V, Kut E, Giotis ES, Skinner MA, et al. Chicken cGAS Senses Fowlpox Virus Infection and Regulates Macrophage Effector Functions. Front Immunol 2020; 11: 613079.

29. Sun L, Wu J, Du F, Chen X, Chen ZJ. Cyclic GMP-AMP synthase is a cytosolic DNA sensor that activates the type I interferon pathway. Science 2013; 339: 786–791.

30. Vanpouille-Box C, Alard A, Aryankalayil MJ, Sarfraz Y, Diamond JM, Schneider RJ, et al. DNA exonuclease Trex1 regulates radiotherapy-induced tumour immunogenicity. Nat Commun 2017; 8: 15618.

31. Yang H, Wang H, Ren J, Chen Q, Chen ZJ. cGAS is essential for cellular senescence. Proc Natl Acad Sci U S A 2017; 114: E4612–E4620.

32. Krausgruber T, Blazek K, Smallie T, Alzabin S, Lockstone H, Sahgal N, et al. IRF5 promotes inflammatory macrophage polarization and TH1-TH17 responses. Nat Immunol 2011; 12: 231–238.

33. Bailey JD, Diotallevi M, Nicol T, McNeill E, Shaw A, Chuaiphichai S, et al. Nitric Oxide Modulates Metabolic Remodeling in Inflammatory Macrophages through TCA Cycle Regulation and Itaconate Accumulation. Cell Rep 2019; 28: 218–230 e217.

34. Gao Z, Li Y, Wang F, Huang T, Fan K, Zhang Y, et al. Mitochondrial dynamics controls anti-tumour innate immunity by regulating CHIP-IRF1 axis stability. Nat Commun 2017; 8: 1805.

35. Bleazard W, McCaffery JM, King EJ, Bale S, Mozdy A, Tieu Q, et al. The dynamin-related GTPase Dnm1 regulates mitochondrial fission in yeast. Nat Cell Biol 1999; 1: 298–304.

36. Chang CR, Blackstone C. Dynamic regulation of mitochondrial fission through modification of the dynamin-related protein Drp1. Ann N Y Acad Sci 2010; 1201: 34–39.

37. Nakahira K, Haspel JA, Rathinam VA, Lee SJ, Dolinay T, Lam HC, et al. Autophagy proteins regulate innate immune responses by inhibiting the release of mitochondrial DNA mediated by the NALP3 inflammasome. Nat Immunol 2011; 12: 222–230.

38. Ye Z, Shi Y, Lees-Miller SP, Tainer JA. Function and Molecular Mechanism of the DNA Damage Response in Immunity and Cancer Immunotherapy. Front Immunol 2021; 12: 797880.

39. Chen Y, Corriden R, Inoue Y, Yip L, Hashiguchi N, Zinkernagel A, et al. ATP release guides neutrophil chemotaxis via P2Y2 and A3 receptors. Science 2006; 314: 1792–1795.

40. Jeong JY, Kim J, Kim B, Kim J, Shin Y, Kim J, et al. IL-1ra Secreted by ATP-Induced P2Y2 Negatively Regulates MUC5AC Overproduction via PLCbeta3 during Airway Inflammation. Mediators Inflamm 2016; 2016: 7984853.

41. Cao DJ, Schiattarella GG, Villalobos E, Jiang N, May HI, Li T, et al. Cytosolic DNA Sensing Promotes Macrophage Transformation and Governs Myocardial Ischemic Injury. Circulation 2018; 137: 2613–2634.

42. Wu Q, Leng X, Zhang Q, Zhu YZ, Zhou R, Liu Y, et al. IRF3 activates RB to authorize cGAS-STING-induced senescence and mitigate liver fibrosis. Sci Adv 2024; 10: eadj2102.

43. Kapetanovic R, Afroz SF, Ramnath D, Lawrence GM, Okada T, Curson JE, et al. Lipopolysaccharide promotes Drp1-dependent mitochondrial fission and associated inflammatory responses in macrophages. Immunol Cell Biol 2020; 98: 528–539.

44. Zaja I, Bai X, Liu Y, Kikuchi C, Dosenovic S, Yan Y, et al. Cdk1, PKCdelta and calcineurin-mediated Drp1 pathway contributes to mitochondrial fission-induced cardiomyocyte death. Biochem Biophys Res Commun 2014; 453: 710–721.

45. Xie Q, Wu Q, Horbinski CM, Flavahan WA, Yang K, Zhou W, et al. Mitochondrial control by DRP1 in brain tumor initiating cells. Nat Neurosci 2015; 18: 501–510.

46. Xu S, Wang P, Zhang H, Gong G, Gutierrez Cortes N, Zhu W, et al. CaMKII induces permeability transition through Drp1 phosphorylation during chronic beta-AR stimulation. Nat Commun 2016; 7: 13189.

47. Kashatus JA, Nascimento A, Myers LJ, Sher A, Byrne FL, Hoehn KL, et al. Erk2 phosphorylation of Drp1 promotes mitochondrial fission and MAPK-driven tumor growth. Mol Cell 2015; 57: 537–551.

48. Han H, Tan J, Wang R, Wan H, He Y, Yan X, et al. PINK1 phosphorylates Drp1(S616) to regulate mitophagy-independent mitochondrial dynamics. EMBO Rep 2020; 21: e48686.

49. Shen X, Sun C, Cheng Y, Ma D, Sun Y, Lin Y, et al. cGAS Mediates Inflammation by Polarizing Macrophages to M1 Phenotype via the mTORC1 Pathway. J Immunol 2023; 210: 1098–1107.

50. Wang Q, Bergholz JS, Ding L, Lin Z, Kabraji SK, Hughes ME, et al. STING agonism reprograms tumor-associated macrophages and overcomes resistance to PARP inhibition in BRCA1-deficient models of breast cancer. Nat Commun 2022; 13: 3022.

51. Ishikawa H, Barber GN. STING is an endoplasmic reticulum adaptor that facilitates innate immune signalling. Nature 2008; 455: 674–678.

52. Hartlova A, Erttmann SF, Raffi FA, Schmalz AM, Resch U, Anugula S, et al. DNA damage primes the type I interferon system via the cytosolic DNA sensor STING to promote anti- microbial innate immunity. Immunity 2015; 42: 332–343.

53. Sarode P, Zheng X, Giotopoulou GA, Weigert A, Kuenne C, Gunther S, et al. Reprogramming of tumor-associated macrophages by targeting beta-catenin/FOSL2/ARID5A signaling: A potential treatment of lung cancer. Sci Adv 2020; 6: eaaz6105.

54. Kaneda MM, Messer KS, Ralainirina N, Li H, Leem CJ, Gorjestani S, et al. PI3Kgamma is a molecular switch that controls immune suppression. Nature 2016; 539: 437–442.

55. Pello OM, Chevre R, Laoui D, De Juan A, Lolo F, Andres-Manzano MJ, et al. In vivo inhibition of c-MYC in myeloid cells impairs tumor-associated macrophage maturation and pro-tumoral activities. PLoS One 2012; 7: e45399.

56. Zhang JL, Liu Y, Yang H, Zhang HQ, Tian XX, Fang WG. ATP-P2Y2-beta-catenin axis promotes cell invasion in breast cancer cells. Cancer Sci 2017; 108: 1318–1327.

57. Hu LP, Zhang XX, Jiang SH, Tao LY, Li Q, Zhu LL, et al. Targeting Purinergic Receptor P2Y2 Prevents the Growth of Pancreatic Ductal Adenocarcinoma by Inhibiting Cancer Cell Glycolysis. Clin Cancer Res 2019; 25: 1318–1330.

58. Katz S, Ayala V, Santillan G, Boland R. Activation of the PI3K/Akt signaling pathway through P2Y(2) receptors by extracellular ATP is involved in osteoblastic cell proliferation. Arch Biochem Biophys 2011; 513: 144–152.

59. Mehta AK, Cheney EM, Hartl CA, Pantelidou C, Oliwa M, Castrillon JA, et al. Targeting immunosuppressive macrophages overcomes PARP inhibitor resistance in BRCA1-associated triple-negative breast cancer. Nat Cancer 2021; 2: 66–82.

60. Zhang F, Parayath NN, Ene CI, Stephan SB, Koehne AL, Coon ME, et al. Genetic programming of macrophages to perform anti-tumor functions using targeted mRNA nanocarriers. Nat Commun 2019; 10: 3974.

61. Allouch A, Voisin L, Zhang Y, Raza SQ, Lecluse Y, Calvo J, et al. CDKN1A is a target for phagocytosis-mediated cellular immunotherapy in acute leukemia. Nat Commun 2022; 13: 6739.

62. Paoletti A, Allouch A, Caillet M, Saidi H, Subra F, Nardacci R, et al. HIV-1 Envelope Overcomes NLRP3-Mediated Inhibition of F-Actin Polymerization for Viral Entry. Cell Rep 2019; 28: 3381–3394 e3387.

63. Allouch A, Di Primio C, Paoletti A, Le-Bury G, Subra F, Quercioli V, et al. SUGT1 controls susceptibility to HIV-1 infection by stabilizing microtubule plus-ends. Cell Death Differ 2020; 27: 3243–3257.

