## Supplementary Information for "P2Y2 purinergic receptor and DNA sensor cGAS dictate ionizing radiation-mediated proinflammatory macrophage activation"

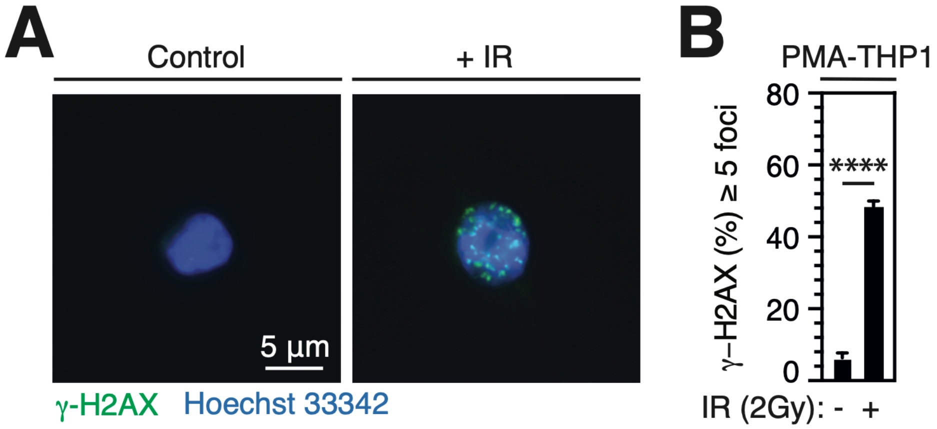


**Supplementary Figure S1: PMA-THP1 macrophages exhibit γ-H2AX$^+$ foci after 2-Gy irradiation. A, B** Fluorescence microscopy images (**A**) and percentages (**B**) of PMA-differentiated human THP1 macrophages showing γ-H2AX$^+$ nuclear foci 1 hour after treatment with control or 2 Gy single-dose IR are shown (scale bar, 5 µm). Nuclei are stained with Hoechst 33342. Data are means ± S.E.M from three independent experiments. P-values (****$P < 0.0001$) were determined with two-tailed unpaired t test (**B**).

**Supplementary Table S1. Human siRNA, siGENOME SMARTpool, siON-TARGETplus SMARTpool and shRNA used in this study.**

| Human Target | Sg | Target sequence | Reference |
|---|---|---|---|
| siGENOME Non Targeting SMARTpool | 1<br>2<br>3<br>4 | UAAGGCUAUGAAGAGAUAC<br>AUGUAUUGGCCUGUAUUAG<br>AUGAACGUGAAUUGCUCAA<br>UGGUUUACAUGUCGACUAA | Dharmacon D-001206-14-05 |
| siGENOME SMARTpool MB21D1 | 1<br>2<br>3<br>4 | GAAGAAACAUGGCGGCUAU<br>GAAGAGAAAUGUUGCAGGA<br>GUAAGGAAUUUCUGACAAA<br>CAACACUCGUGCAUAUUAC | Dharmacon M-015607-01-0005 |
| siON-TARGET plus Non Targeting SMARTpool | 1<br>2<br>3<br>4 | UGGUUUACAUGUCGACUAA<br>UGGUUUACAUGUUGUGUGA<br>UGGUUUACAUGUUUUCUGA<br>UGGUUUACAUGUUUUCCUA | Dharmacon D001810-10-05 |
| siON-TARGET plus SMARTpool DNM1L | 1<br>2<br>3<br>4 | GGAGCCAGCUAGAUAUUAA<br>CAUCAGAGAUUGUUUACCA<br>CGUAAAAGGUUGCCUGUUA<br>GUGAACCCGUGGAUGAUAA | Dharmacon L-012092-00-0005 |
| siON-TARGET plus SMARTpool P2Y2 (siP2Y2.3) | 1<br>2<br>3<br>4 | CGAGAACACUAAGGACAUU<br>CGACAGAACUGACAUGCAG<br>GGAAUGCGUCCACCACAUA<br>UGCCGCUGCUGGUCUAUUA | Dharmacon L-003688-00-0005 |
| siControl | 1 | UUCAAUAAAUUCUUGAGGU | Sigma-Aldrich |
| siP2Y2.1 | 1 | GGAUAGAAGAUGUGUUGGG | Sigma-Aldrich |
| siP2Y2.2 | 2 | GGCUGUAACUUAUACUAAA | Sigma-Aldrich |
| pLKO 1 shP2Y2 | 1 | ATGTTCCACCTGGCTGTGTCTGATGCACT | Sigma-Aldrich |
